# Expectancy changes the self-monitoring of voice identity

**DOI:** 10.1101/2020.07.22.215350

**Authors:** Joseph F. Johnson, Michel Belyk, Michael Schwartze, Ana P. Pinheiro, Sonja A. Kotz

## Abstract

Self-voice attribution can become difficult when voice characteristics are ambiguous, and functional magnetic resonance imagines (fMRI) investigations of such ambiguity are sparse. We utilized voice-morphing (self-other) to manipulate (un-)certainty in self-voice attribution in a button-press paradigm. This allowed investigating how levels of self-voice certainty alter brain activation in regions monitoring voice identity areas and unexpected changes in voice playback quality. FMRI results confirm a self-voice suppression effect in the right anterior superior temporal gyrus (aSTG) when self-voice attribution was unambiguous. Although the right inferior frontal gyrus (IFG) was more active during self-generated voice compared to when passively-heard, the putative role of this region in detecting unexpected self-voice changes was not confirmed. Further research on the link between right aSTG and IFG is required and may establish a threshold monitoring voice identity in action. The current results have implications for a better understanding of an altered experience of self-voice feedback leading to auditory verbal hallucinations.

## INTRODUCTION

Self-monitoring of the voice relies on comparing what we expect to hear and what we actually hear (Frith, 1992; Wolpert et al., 1998). However, in a dynamic environment sensory feedback is often ambiguous, e.g., when listening to multiple speakers. Any judgment of the voice source further depends on how much sensory feedback deviates from expectations (Feinberg, 1978). Minor deviations regarding one’s own voice are typically self-attributed and used to compensate motor control. Major deviations may lead to source-attributing the voice to another person. People who experience auditory verbal hallucinations (AVH) show dysfunctional self-monitoring (Kumari et al., 2010b; Sapara et al., 2015). For example, schizophrenia patients who experience AVH are more likely to incorrectly attribute their voice to an external source in ambiguous conditions that result in uncertainty among healthy individuals (Johns et al., 2001; Allen et al., 2004; Pinheiro et al., 2016a). However, AVH are not limited to persons with psychosis but are also experienced along a spectrum of hallucination proneness in healthy individuals (Baumeister et al., 2017). An externalization bias observed within the general population may relate to higher proneness to experience AVH in otherwise healthy individuals (Asai & Tanno, 2013; Pinheiro et al., 2019). Functional neuroimaging studies of self-voice monitoring in the healthy brain have examined the neural substrates of self-other voice attribution but have so far not examined responses to uncertainty in ambiguous conditions (e.g. Allen et al., 2006; Fu et al., 2006). It is critical to not only know how the brain establishes correct self and other voice attribution but also where and how the voice is processed in conditions of uncertainty to gain a better understanding of the mechanisms underlying dysfunctional self-monitoring.

Previous research has reported that unaltered self-voice production leads to reduced functional brain activity in the auditory cortex (Christoffels, Formisano, & Schiller, 2007). This motor-induced suppression (MIS) is compatible with findings of numerous studies employing diverse methodology. It is similar to the N1 suppression effect, a modulation of the event-related potential of the electroencephalogram (EEG) (e.g. Heinks-Maldonado, Mathalon, Gray, & Ford, 2005; Behroozmand & Larson, 2011; Sitek et al., 2013; Wang et al., 2014; Pinheiro, Schwartze, & Kotz, 2018), or M1 suppression in magnetoencephalography (Numminen, Salmelin, & Hari, 1999; Houde et al., 2002; Ventura, Nagarajan, & Houde, 2009), weakened activity in electrocorticography and at intracranial electrodes (Greenlee et al.,2011; Chang et al., 2013), or direct- and inter-cell recordings in non-human primates (Müller-Preuss & Ploog, 1981; Eliades & Wang, 2008). Contrasting with the suppressed activity in auditory cortex, self-voice monitoring activates a widespread system of functionally connected brain regions, including the inferior frontal gyrus (IFG), supplementary motor area, insula, pre- and postcentral gyrus, inferior parietal lobule (IPL), motor cortex, thalamus, and cerebellum (Christoffels, Formisano, & Schiller, 2007; Behroozmand et al., 2015). The right anterior superior temporal gyrus (aSTG) and the adjacent upper bank of the superior temporal sulcus (STS) likely play a critical role in voice identity perception (Belin, Fecteau, & Bedard, 2004; von Kriegstein et al., 2003; von Kriegstein & Giraud, 2004; Belin and Zatorre, 2003). Patient studies support this assumption as lesions or damage to the aSTG can lead to deficits in voice identity recognition (Gainotti, Ferraccioli, & Marra, 2010; Gainotti & Marra, 2011; Hailstone et al., 2011; van Lancker & Kreiman, 1987; van Lancker & Canter, 1982).

MIS in voice monitoring is not only effective in voice production but also in response to voice recordings activated via a button press (Ford et al., 2007; Whitford et al., 2011; Pinheiro, Schwartze, & Kotz, 2018; Knolle, Schwartze, Schröger, & Kotz, 2019) as well as for non-verbal sounds including tones (e.g. Aliu, Houde, & Nagarajan, 2009; Baess, Widmann, Roye, Schröger, & Jacobsen, 2009; Knolle, Schröger, & Kotz, 2013). Moreover, MIS seems to operate across modalities of sensory feedback and arises from various motor effectors (e.g. Miall & Wolpert, 1996; Wolpert et al., 1998; Leube et al., 2003; Blakemore, Wolpert, & Frith, 1998). One explanation for MIS is that internal models of expected action outcomes are fed-forward to the relevant cortical regions to cancel out impending activity to the anticipated stimulus (Jordan & Rumelhart, 1992; Miall & Wolpert, 1996; Wolpert, 1997). Studies that experimentally manipulate sensory feedback create a mismatch between expected and actual outcome and indicate concomitant modulation or absence of MIS under such circumstances. EEG studies typically show this as decreased N1 suppression (e.g. Heinks-Maldonado et al., 2005; Behroozmand & Larson, 2011), while fMRI studies report a relative increase of STG activity when expected feedback is altered (McGuire, Silbersweig, & Frith, 1996; Fu et al., 2006; Christoffels et al., 2007; Zheng et al., 2010; Christoffels et al., 2011). With this approach, it is not only possible to make listeners uncertain about self- or other-voice attribution (Allen et al., 2004, 2005, 2006; Fu et al., 2006; Vermissen et al., 2007), but to also lead listeners to incorrectly attribute self-voice to another speaker (Johns et al., 2001, 2003, 2006; Fu et al., 2006; Allen et al., 2004, 2005, 2006, Kumari et al., 2010a, 2010b; Sapara et al., 2015). STG suppression only persists when the voice is correctly judged as self-voice in distorted feedback conditions (Fu et al., 2006). Critically, data reflecting uncertain voice attribution are often removed from fMRI analyses (Allen et al., 2005; Fu et al., 2006). However, in order to gain a better understanding of voice attribution to internal or external sources, it is mandatory to specifically focus on such data and to define how the known voice attribution region of the STG reacts in conditions of uncertainty.

In addition to auditory cortex, activation in the right inferior frontal gyrus increases in response to distorted auditory feedback (Johnson et al., 2019). However, while attenuation of the right aSTG activation reflects expected stimulus quality, the right IFG is selectively responsive to unexpected sensory events (Aron, Robbins, & Poldrack, 2004). Increased right IFG activity has been reported when feedback is acoustically altered (Behroozmand et al., 2015, Fu et al., 2006; Toyomura et al., 2007; Tourville et al., 2008; Guo et al., 2016), delayed (Sakai et al., 2009; Watkins et al., 2005), replaced with the voice of another speaker (Fu et al., 2006), or physically perturbed during vocal production (Golfinopoulos et al., 2010). In response to unexpected sensory feedback in voice production, the right IFG produces a “salient signal”, indicating the potential need to stop and respond to stimuli that may be affected by or the result of some external influence. Correspondingly, It has been hypothesized that the processing of salient stimuli with minimal divergence from expectations leads to an externalization bias that may manifest in the experience of AVH (Sommer et al., 2008).

In the current fMRI experiment, we investigated how cortical voice identity and auditory feedback monitoring regions respond in (un)certain self-other voice attribution. Participants elicited voice stimuli that varied along a morphing continuum from self to other voice, including intermediate voice samples of ambiguous identity. Region of interest (ROI) analyses motivated by our research question and a priori hypotheses focussed on the right aSTG and the right IFG. The right aSTG ROI stems from a well-replicated temporal voice area (TVA) localizer task (Belin et al., 2000). The right IFG ROI conforms to a region responsive to experimental manipulation of auditory feedback previously identified via activation-likelihood estimation (ALE) analyses (Johnson et al., 2019). Due to possible individual variability in thresholds for self-other voice attribution (Asai & Tanno, 2013), each participant underwent psychometric testing to determine individualized points of maximum uncertainty on a continuum from self to other voice. The primary goal was to test hypotheses that i) MIS of self-voice in the right aSTG is present, and the degree of suppression is greater when attribution of the self is certain compared to when uncertain, and ii) right IFG activation would increase in response to an increase in voice uncertainty. Confirmation of these results would further substantiate previous EEG findings regarding MIS for self-voice elicited via button-press as compared to passively heard (Ford et al., 2007; Whitford et al., 2011; Pinheiro, Schwartze, & Kotz, 2018; Knolle, Schwartze, Schröger, & Kotz, 2019), indicating that suppressed activity in auditory cortex aligns with predicted self-voice quality and not only as a function of expected quality of voice feedback.

## METHODS

### PARTICIPANT RECRUITMENT

Twenty-seven participants took part in the study. The data of two participants were discarded due to scanning artefacts. Of the remaining 25 (17 female), the average age was 21.88 years (SD = 4.37; range 18 to 33). Inclusion criteria assured that participants had no diagnosis of psychological disorder, normal or corrected-to-normal hearing and vision, and no evidence of phonagnosia. This was tested using an adapted version of a voice-name recognition test described below (Roswandowitz et al., 2014). All participants gave informed consent and were compensated with university study participant credit. This study was approved by the Ethical Review Committee of the Faculty of Psychology and Neuroscience at Maastricht University (ERCPN-176_08_02_2017).

### PROCEDURES

#### PHONAGNOSIA SCREENING

Phonagnosia is a disorder restricting individuals from perceiving speaker identity in voice (Van Lancker et al., 1988). We screened for phonagnosia using an adapted version of a phonagnosia screening task (see Roswandowitz et al., 2014). The task was composed of four rounds of successive learning and testing phases, in which participants initially listened to the voices of three speakers of the same gender. Identification of each speaker was subsequently tested 10 times with response accuracy feedback provided during the first half of test trials. Finally, the task was repeated with stimuli of the gender not used in the first run. Presentation order of these runs was counterbalanced across participants.

#### PSYCHOMETRIC TASK

In a voice attribution task (VAT), participants heard samples of the vowels /a/ and /o/. These samples varied in voice identity, which was morphed along a continuum from “self-voice” to “other-voice” using the STRAIGHT voice morphing software package (Kawahara 2003, 2006) running in MATLAB (R2019A, v9.6.0.1072779, The MathWorks, Inc., Natick, MA). For this procedure, samples of the self-voice (SV) and other voice (OV) producing the two vowels were obtained from each participant and normalized in duration (500ms) and amplitude (70db) using the Praat software package (v6.0.28, http://www.praat.org/). The OV sample used matched the gender of the participant. On this basis, 11 stimuli were created along a morphing spectrum in steps of 10% morphing from SV to OV. In a two-alternative forced-choice (2AFC) task, participants listened to each stimulus presented in random order and responded to the question: Is the voice “more me” or “more other”? This procedure was repeated twice. In one run stimuli were presented passively while in the other run participants were visually cued to press a button which elicited the next stimulus (see Figure 1). This procedure was used to identify an individualized point of maximum ambiguity (PMA) along the morphing spectrum for each participant. The PMA was defined as the stimulus that was closest to chance level (50%) and used to inform subsequent fMRI analyses.

**Figure 1.**
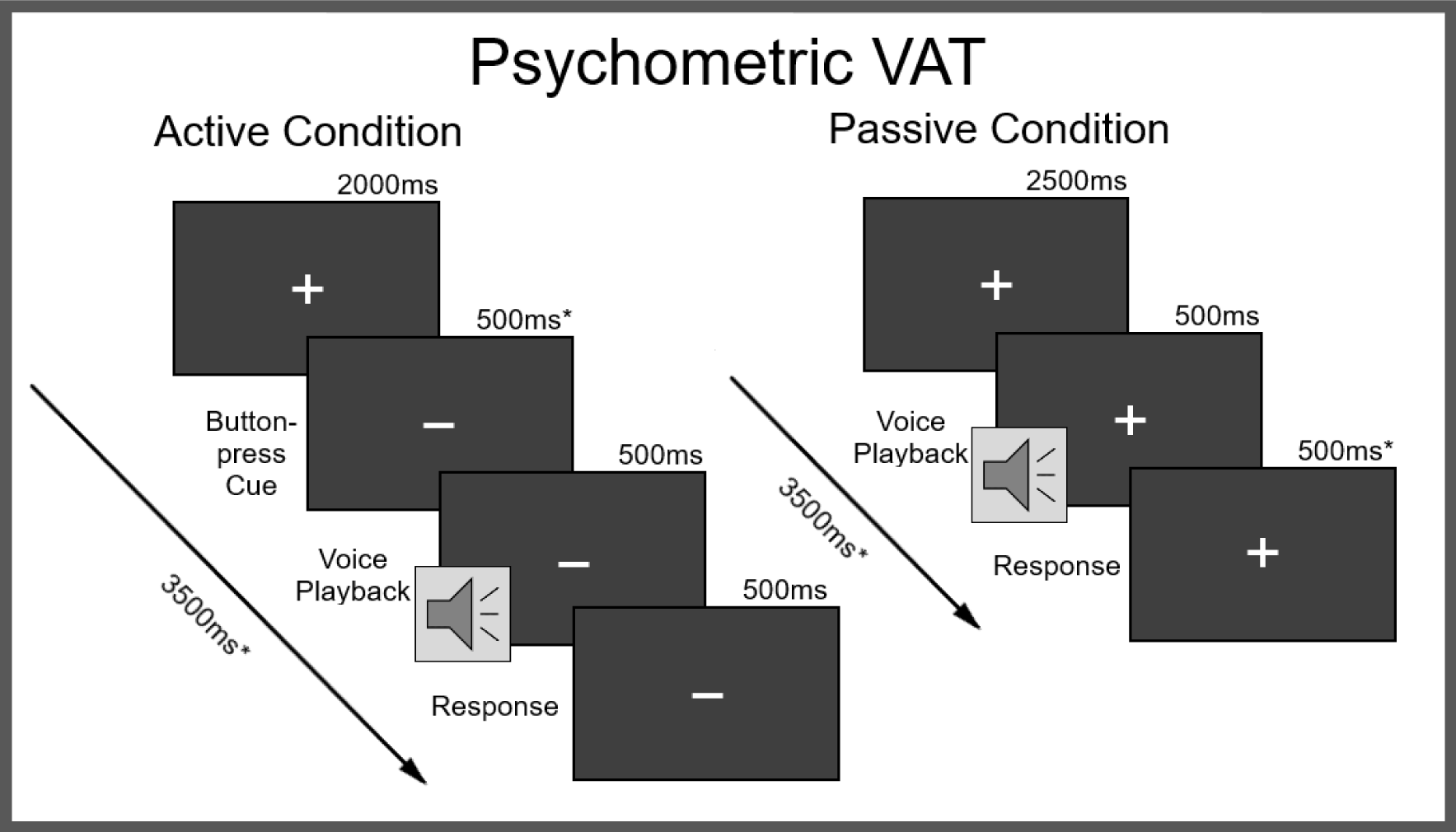
Psychometric Voice Attribution Task (VAT): Active = button-press condition; Passive = hearing conditions, * = affected by individual motor response-time variability; Response = two-alternate forced-choice (“The voice sounded more like me.” or “The voice sounded more like someone else.”).

#### FMRI TASKS

##### Temporal Voice Area (TVA) Localizer

To identify voice sensitive brain areas, participants were scanned during a voice localizer task (Belin et al., 2000). This task is widely used to reliably probe activity along the bilateral temporal cortices (e.g. Pernet et al., 2015) designated as anterior, middle, and posterior TVA regions. Stimuli consisted of 8-second auditory clips with 20 vocal and 20 non-vocal sounds. In a single run, participants passively listened to these sounds and 20 silent trials in pseudorandom order. Contrasting responses to vocal and non-vocal sounds identified brain regions selectively sensitive to voice processing. The peak activation in the anterior STG of the right hemisphere was then chosen as the voice-attribution ROIs in the subsequent fMRI analysis.

##### Voice Perception Task (VPT)

Participants listened to passively presented or self-generated voice stimuli. When shown a cue signifying the active button-press condition, participants pressed a button to elicit voice stimuli, and conversely when shown a cue signifying the passive condition were instructed to do nothing (Figure 2). In the active condition, half of the trials elicited a voice following the button press, while in the other half no voice was presented. In the passive condition, all trials involved the presentation of a voice. A subset of stimuli used in the VAT was selected for the VPT, specifically the 100, 60, 50, 40, and 0% self-voice morphs. The intermediate steps of 60, 50, and 40% were selected as piloting revealed that individual PMA fell within a range of 35-65% morphing, while morphs outside of this range produced high degrees of confidence in self vs. other judgement. This ensured that every participant received the voice stimuli nearest to their subjective PMA. Trial onsets were 9 seconds (+/-500ms) apart to allow the BOLD response to return to baseline before the presentation of the next stimulus started. To avoid the effects of adaptation suppression (Andics et al., 2010, 2013; Belin et al., 2003; Latinus & Belin., 2011; Wong et al., 2004), voice conditions were presented in a random order. Over two runs, a total of 100 trials were presented in each condition of Source (Active and Passive). Within each condition of Source, each voice stimulus (100, 60, 50, 40, and 0% morphs from self-to-other) was heard 20 times. 20 null trials were included to provide a baseline comparison of activity in response to experimental trials.

**Figure 2.**
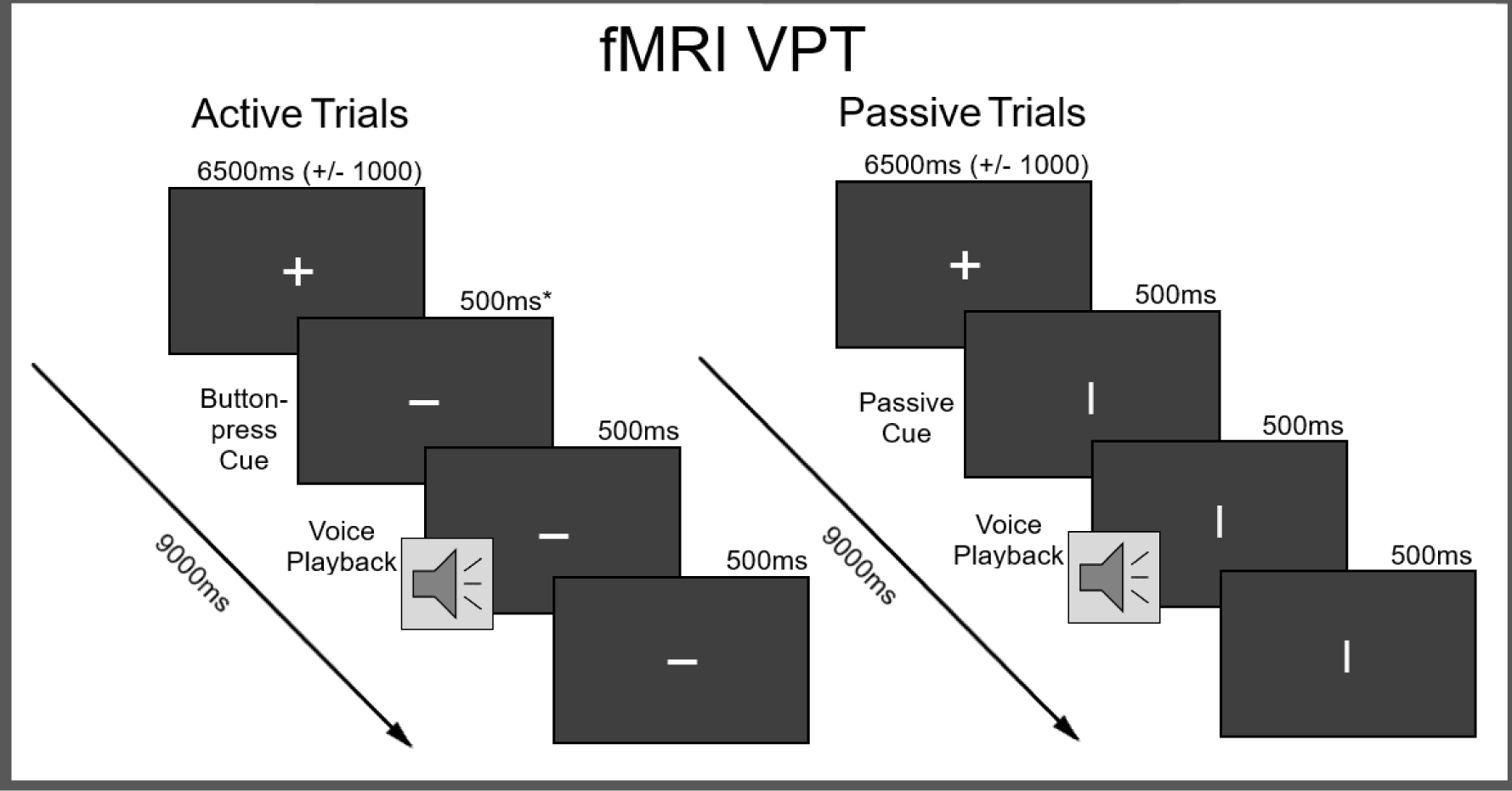
fMRI Voice Perception Task (VPT): Active = button-press condition; Passive = hearing conditions, * = affected by individual motor response-time variability.

### FMRI DATA ACQUISITION AND ANALYSIS

Data acquisition was performed at a Siemens 3T Magnetom Prisma Fit Magnetic Resonance Imaging (MRI) scanner at Scannexus facilities (Maastricht, NE), equipped with a 32-channel head coil (Siemens Healthcare, Erlangen, Germany). A structural whole brain T1-weighted single-shot echoplanar imaging (EPI) sequence was collected for each participant (field of view (FOV) 256mm; 192 axial slices; 1mm slice thickness; 1 x 1 x 1mm voxel size; repetition time (TR) of 2250ms seconds; echo-time (TE) 2.21ms). Two functional tasks were conducted with T2-weighted EPI scans (FOV 208mm; 60 axial slices; 2mm slice thickness; 2 x 2 x 2mm voxel size; TE 30ms; flip angle = 77°). Both tasks applied a long inter-acquisition interval where time between consecutive image acquisition (2000ms) was delayed, resulting in a TR of 10 and 9 seconds for the TVA localizer and VPT, respectively. This allowed auditory stimuli to be presented during a period of relative silence to reduce noise artifacts and for volume acquisition to proceed during a period of peak activation in the auditory cortex (Belin et al., 1999; Hall et al., 1999).

DICOM image data was converted to 4D NIFTI format using the Dcm2Nii converter provided in the MRIcron software package (https://www.nitrc.org/projects/mricron/) www.nitrc.org/projects/mricron/). The topup tool (Smith, et al., 2004) implemented in FSL (www.fmrib.ox.ac.uk/fsl) was used to estimate and correct for susceptibility induced image distortions. Pre-processing was performed using SPM12 (Wellcome Department of Cognitive Neurology, London, UK). A pre-processing pipeline applied slice timing correction, realignment and unwarping, segmentation, normalization to standard Montreal Neurological Institute (MNI) space (Fonov et al., 2009) as well as smoothing with a full width at half maximum (FWHM) 8mm isotropic Gaussian kernel.

#### General Linear Model (GLM) Analysis

The TVA localizer and experimental VPT fMRI data were analyzed with a standard two-level procedure in SPM12. For the TVA localizer, contrast images for Vocal > Non-Vocal and Vocal > Silent were estimated for each participant. To test for the main effect of interest, conjunction analysis ((V > NV) ∩ (V > S)) was performed. A second level random-effects analysis tested for group-level significance. A first-level fixed-effects GLM of the VPT data calculated contrast estimates for each participant. Contrast estimates were then used in the subsequent hypothesis-driven ROI analysis to investigate TVA activity.

#### Linear Mixed Model (LMM) ROI Analyses

Two spherical (5mm) ROIs were selected for analysis: the right aSTG/S in Brodmann Area (BA) 22 (MNI coordinates × 58, y 2, z -10) defined by the TVA fMRI localizer task, and the right IFG opercular region in BA 44 (MNI coordinates x 46, y 10, z 4) (See Figure 3). A 2×3 factorial design was formulated using the factors of Source and Voice. The two-leveled factor Source included self-generated (Active: A) and passively-heard (Passive: P) playback of voice recordings. The three-leveled factor Voice included self-identified voice (Self-voice: SV), externally-identified voice (Other-voice: OV), and voice of ambiguous identity (Uncertain: UV) unattributed to self or external.

**Figure 3.**
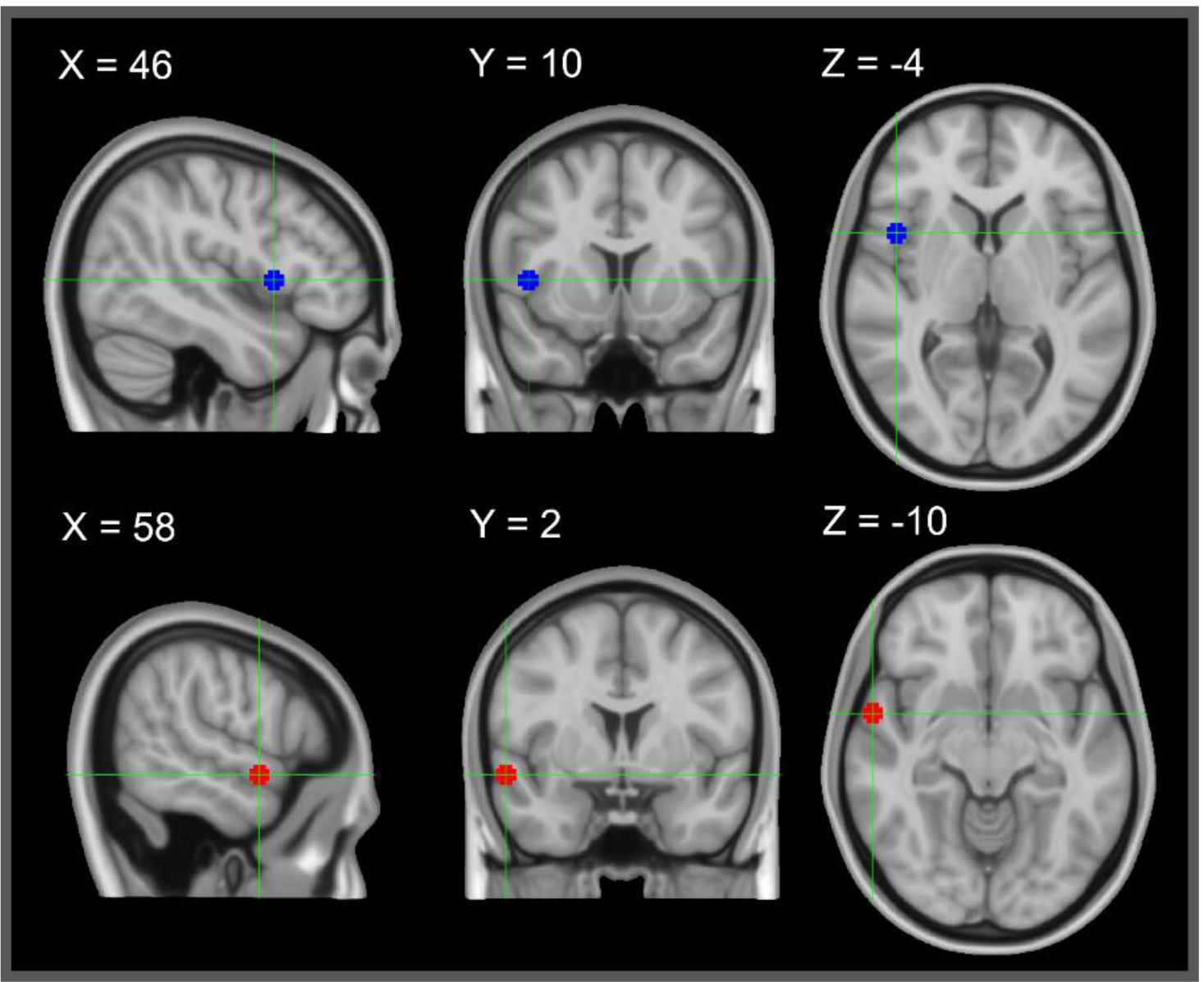
fMRI Regions of Interest: Blue: inferior frontal gyrus; MNI coordinates x 58, y 2, z -10; determined from ALE neuroimaging meta-analysis (Johnson et al., 2019). Red: right anterior superior temporal gyrus; MNI coordinates x 46, y 10, z 4; determined from fMRI temporal voice area localizer task.

Data were analyzed in R v3.6.1 (R Core Team, 2019) running on OS v10.11.6. Data handling and visualization were supplemented with the tidyverse (Wickham, 2017). Linear Mixed Models (LMMs) were fit with lme4 (Bates, Maechler, Bolker, & Walker, 2015). Separate LMMs were fitted for contrast estimates of the IFG and the aSTG ROIs with Source (A and P), Voice (SV, OV and UV), and their interaction as fixed effects. Participant was modelled as a random intercept. Model residuals were examined for potential outliers. Five data points were removed from the IFG analysis and one was removed from the aSTG analysis.

The main effects of Voice, Source and their interaction were tested with the afex package using Kenward-Rogers degrees of freedom (Singmann et al., 2015). Estimated marginal means and confidence intervals were computed with the emmeans package (Lenth, 2020) for visualization. All p-values are corrected for multiple comparisons controlling at a false-discovery rate (FDR) of 0.05.

### VAT RESULTS

Psychometric analysis of the VAT indicated little variability in the degree of morphing between SV and OV required to elicit responses at chance level (50%), which we identified as the point of maximum ambiguity. For the A condition, nine participants had PMAs at 40%, eight at 50% and ten at 60% morphing. In the passive condition, eleven required 40%, seven 50%, and nine 60% morphing. There was no significant difference between the average morphing required to elicit PMA in A (μ 50%, SD 0.085) and P (μ 50%, SD 0.087) conditions. Although no participant matched criteria for phonagnosia as specified by the screening task, VAT data from one participant was excluded due to an inability to reliably differentiate between their own voice and other voices.

### TVA LOCALIZER RESULTS

The TVA fMRI localizer produced four significant cluster-level activations (see Table 1 for details). Within two large bilateral STG (BA 22) clusters, each included three peak-level significant activations. These peaks correspond to the posterior (pSTG), middle (mSTG), and aSTG. Two smaller clusters were found in the right precentral gyrus (BA 6), the left IFG (BA 44), and the left IPL (BA 40). All significant cluster- and peak-level coordinates reported survived an FDR correction of 0.05. The right aSTG peak was chosen for ROI analyses as the voice-attribution ROI. These results replicate the pattern of TVA regions of peak activity (e.g. Belin et al., 2000; Fecteau et al., 2004; Latinus et al., 2013; Pernet et al., 2015).

**Table 1.** TVA Localizer Results. Results from TVA localizer task: Coordinates listed in MNI space; (p/a/m)STG: posterior/anterior/middle superior temporal gyrus, preCG: precentral gyrus, IFG: inferior frontal gyrus; 7 peak-level activations in 4 clusters: 1. left STG, 2. right STG, 3. right preCG, 4. left IFG; All listed significant regions survived FDR-corrected threshold 0.05.

**1. TVA Localizer Results**
| Cluster # | Peak Label | BA | Coordinates (x, y, x) | | | Cluster-Level $p$-FDR | Peak-Level $p$-FDR | Cluster Size (voxels) |
| --- | --- | --- | --- | --- | --- | --- | --- | --- |
| 1 | L pSTG | 22 | -60 | -24 | 0 | $2.67 \times 10^{-14}$ | $9.29 \times 10^{-12}$ | 4551 |
| | L aSTG | 22 | -58 | -10 | -2 | | $9.04 \times 10^{-11}$ | |
| | L mSTG | 22 | -66 | -16 | -2 | | $3.51 \times 10^{-8}$ | |
| 2 | R pSTG | 22 | 58 | -24 | -2 | $2.05 \times 10^{-14}$ | $1.07 \times 10^{-9}$ | 4565 |
| | R aSTG | 22 | 58 | 2 | -10 | | $2.00 \times 10^{-9}$ | |
| | R mSTG | 22 | 58 | -8 | -6 | | $2.74 \times 10^{-9}$ | |
| 3 | R preCG | 6 | 52 | 52 | 0 | 0.007 | $3.44 \times 10^{-4}$ | 408 |
| 4 | L IFG | 44 | -42 | 14 | 22 | 0.019 | 0.002 | 294 |

### LMM ROI RESULTS

Linear mixed model analysis of the right aSTG (Table 2A, Figure 4A) produced an FDR-corrected significant main effect for the factor of voice, (F (2, 118.94) = 4.90, p = 0.021). No significant effect was observed for source (F (1, 118.92) = 0.53, p = 0.47). A trend for the expected interaction effect between voice and source was observed, although did not survive FDR correction for multiple comparisons (F (2, 118.94)3.40, p = 0.065). However, based on our hypotheses and the observed trend we conducted an exploratory post-hoc analysis to test the hypothesis that the contrast A > P differs for SV stimuli as compared to stimuli with OV or UV identities. This was confirmed by the finding that motor-induced suppression is observed preferentially for SV stimuli (t(119) = -2.7, p = 0.021).

**Figure 4.**
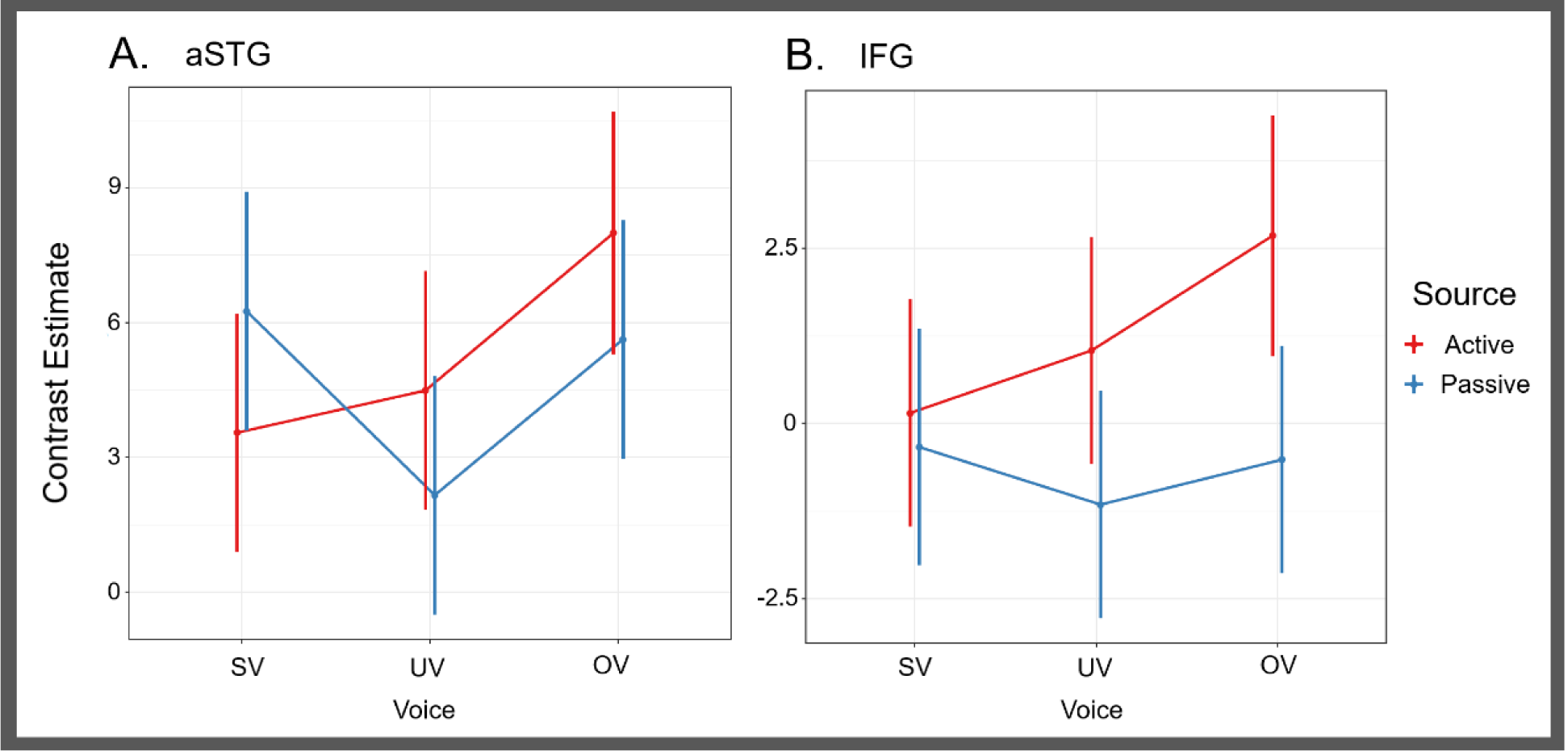
fMRI Voice Perception Task (VPT) LMM Results: Linear mixed model analysis on ROIs in A) right anterior superior temporal gyrus (aSTG) and B) right inferior frontal gyrus (IFG). Active: button-press condition, Passive: passive hearing condition, SV: self-voice, UV: uncertain-voice, OV: other-voice. Post-hoc analysis in right aSTG revealed motor induced suppression (for contrast Active > Passive) for only SV as compared to UV or OV (t(119) = -2.7, p = 0.021).

The LMM analysis was repeated for the right IFG ROI (Table 2B, Figure 4B). A significant FDR-corrected main effect of source was observed (F (1, 116.04) = 9.93, p = 0.002). No main effect was found for the factor of voice (F (2, 115.95) = 1.52, p = 0.26), and no interaction between voice and source were observed (F (2, 115.81) = 1.60, p = 0.26).

## DISCUSSION

The current study investigated the interplay of auditory feedback regions involved in the processing of (un)certainty in self-voice attribution, and unexpected quality of voice feedback. We report the first fMRI evidence congruent with EEG reports that indicates self-voice MIS can be observed even when voice stimuli are elicited by a button press rather than spoken. The predictable qualities learned by long-term experience with self-voice feedback therefore are sufficient to modulate MIS. Importantly, this effect was specific to vocal sounds matching the timbre of the participant’s own voice and was not observed when hearing the voice of another or being uncertain about a speaker. The right IFG pars opercularis showed increased activation in response to self-initiated voice relative to passive exposure. It is plausible that this differential response pattern is driven by the higher proportion of voice trials not attributed to oneself. This region is known to be more active when perceived stimuli are in conflict with expected sensory feedback. Together, these findings suggest a differentiation between and an potential interplay of right IFG and aSTG in voice processing, and more specifically feedback monitoring of self-generated voice and differentiation of self- and other attribution.

### VOICE IDENTITY AND MOTOR-INDUCED SUPRESSION IN THE STG

Our results confirm right aSTG/S involvement in processing voice identity and indicate that it may play a particular role in segregating the speaker’s voice from external voices when monitoring auditory feedback. We replicate previous TVA findings that the STG and upper bank of the STS contain three bilateral voice patches (Table 1) (Belin et al., 2000; Pernet et al., 2015). The processing of speech-related linguistic (“what”) features have been attributed predominantly to the left hemisphere, while speaker-related paralinguistic (“who”) feature processing has been attributed predominantly to the right hemisphere (Belin et al., 2002; Formisano et al., 2008; Moerel, De Martino, & Formisano, 2012; Grandjean et al., 2005; Ethofer et al., 2006; Schirmer & Kotz, 2006; Ethofer et al., 2007; Wiethoff et al., 2008; Kotz et al., 2003). Moreover, right hemisphere paralinguistic processing of speaker-identity has been localized to the anterior region of the STG/S (Belin et al., 2003; von Kriegstein et al., 2003; von Kriegstein & Giraud, 2004; Fecteau et al., 2004; Latinus et al., 2013; Schelenski, Borowiak, & von Kriegstein, 2016). Different low-level acoustics used in voice identity perception are processed in the pSTG, the extracted cues relevant for speaker identification are then processed in the mSTG, and finally differential processing of voice identity occurs in aSTG (Maguiness et al., 2019). To provide sufficient duration for the extraction of paralinguistic speaker-related features, steady 500ms vowel excerpts were chosen as voice samples in our study (Pinheiro et al., 2018; Schweinberger et al., 2011; Schweinberger, Herholz, & Sommer, 1997; Van Berkum, van den Brink, Tesink, Kos, & Hagoort, 2008). Although vowels provide fundamental cues that allow differentiating between speakers (Belin, Fecteau, & Bedard, 2004; Kreiman & Sidtis, 2011; Latinus & Belin, 2011; Schweinberger et al., 2014), to the best of our knowledge no study has yet confirmed whether such basic stimuli carry enough identity cues to allow for explicit self-recognition (Conde, Goncalves, & Pinheiro, 2018). We conducted ROI analyses on voice identity processing only in the right aSTG due to its responsiveness to variation in voice identity and did not include other TVA regions in our analysis. In doing so, this allowed us to detect fine-grain differences in activation patterns influenced by only identity processing in a region that is highly active in the perception of the voice. We confirmed prior to fMRI testing via psychometric analysis on behavioural data the ability for participants to correctly attribute voice to self and other. Furthermore, we provide the first evidence that 500ms steady vowel recordings of SV and OV allow for accurate recognition of self- and other-attribution.

We observed that motor induced suppression in the right aSTG occurred only for SV. One possible interpretation for this selective finding is that participants are most familiar with the acoustic characteristics of their self-voice, and that they can therefore predict the features of their self-voice more efficiently. Activity in cortical sensory processing regions activate more strongly by stimuli that are unexpected than stimuli that are easily predicted. In the right aSTG, voice identity processing is determined by the extent that speaker-related cues deviate from prototypes of expected voice qualities (Mullenix et al., 2011; Bruckert et al., 2010; Andics et al., 2010, 2013; Latinus & Belin, 2011; Latinus et al., 2013; Petkov & Vuong, 2013; Schweinberger et al., 2014). These prototypes are learned through mean-based coding (Hoffman & Logothesis, 2009). While it is clear that low-level acoustic processing is involved in this comparison (Smith & Patterson, 2005; Smith, Waters, & Patterson, 2007; Gaudrain et al., 2009; Baumann & Belin, 2010; Nolan, McDougall, & Hudson, 2011; Zheng et al., 2011; Kreitewolf, Gaudrain, & von Kreigstein, 2014), the specific features which drive identification vary from voice to voice (Lavner, Gath, & Rosenhouse, 2000; Lavner, Rosenhouse, & Gath, 2001; Kreiman et al., 1992; Latinus & Belin, 2012; Xu et al., 2013). Furthermore, variability in the acoustic features of the voice do not only exist between speakers, but also occurs within individual speakers (Lavan et al., 2019). Therefore, increased experience with the voice of a specific speaker facilitates more efficient recognition of voice identity. Indeed, people have the most experience with the qualities of their self-voice, allowing for easy identification of their own identity, as little divergence from mean-based coding is detected.

Alternatively, MIS of self-voice in a dynamic multi-speaker environment is important for the segregation of internally- and externally-controlled voice stimuli. As verbal communication is typically performed with the synchronized perception of one’s own voice, the sound of self-voice may therefore gain a privileged status that is also reflected in auditory feedback processing. During vocalization, an efference copy of the motor command is sent from motor planning areas to auditory and sensorimotor cortical regions to notify of impending feedback (Rauschecker & Scott, 2009; Rauschecker, 2011; Tourville & Guenther, 2011; Hickok, Houde, & Rong, 2011; Hickok, 2012; Kearney & Guenther, 2019). In specific, error-cells in the pSTG (planum temporale) receive these signals from Broca’s to remain inactive in response to the expected sound of self-voice, and to engage when perceiving voice feedback outside the control of the speaker (Guenther et al., 2006). To date, fMRI research using vocal feedback paradigms has provided evidence for this form of MIS dependent on vocal production. For example, MIS has been reported for unaltered vocal production relative to when hearing a recording of self-voice or when in a noisy environmental (Christoffels et al., 2007), when acoustically distorted (McGuire, Silbersweig, & Frith, 1996; Fu et al., 2006; Zheng et al., 2010; Christoffels et al., 2011) or replaced with the voice of another speaker (McGuire, Silbersweig, & Frith, 1996; Fu et al., 2006). However, as these paradigms all rely on vocal production, they are unable to indicate how the identity processing region of the STG responds specifically to self-identity in voice during action. EEG research has provided evidence for MIS in the auditory cortex that does not depend on vocal speech production as it is observed even when sounds are elicited by a button press. For example, MIS of the N1 response was reported for both, vocal (Heinks-Maldonado, Mathalon, Gray, & For, 2005; Behroozmand & Larson, 2011; Sitek et al., 2013; Wang et al., 2014; Pinheiro, Schwartze, & Kotz, 2018) and button-press elicited self-voice (Ford et al., 2007; Whitford et al., 2011; Pinheiro, Schwartze, & Kotz, 2018; Knolle, Schwartze, Schröger, & Kotz, 2019). In line with previous EEG evidence, the current findings of our button-press fMRI experiment in the voice identity auditory cortex ROI (right aSTG) indicate suppressed activity in response to self-attributed voice during action. The reported MIS is specific to self-voice processing, providing further evidence of voice identity suppression separate from previously described cortical suppression during unperturbed speech. Importantly, this pattern was observed only for voice attributed to oneself with certainty, and not present when voice was distorted to an extent where self-attribution was uncertain.

### EXPECTED FEEDBACK AND THE IFG

The right IFG was more strongly activated when participants generated vocal stimuli with a button press as compared to passive perception. This finding confirms that this region is more responsive to sounds triggered by the participant, potentially as part of auditory feedback. Increased activity in this region has been observed in response to acoustically altered (Behroozmand et al., 2015, Fu et al., 2006; Toyomura et al., 2007; Tourville et al., 2008; Guo et al., 2016), physically perturbed (Golfinopoulos et al., 2010), and externalized voice feedback (Fu et al., 2006).

In response to unexpected sensory information, the right IFG plays a crucial role in relaying salient signals to attention networks. Moreover, the right IFG pars opercularis is part of a prediction network, which forms expectations and detects unexpected sensory outcomes (Siman-Tov et al., 2019). When prediction errors are detected, an inferior frontal network produces a salience response (Cai et al., et al., 2014; Seeley, 2010; Power et al., 2011; Chang et al., 2013). Salience signals engage ventral and dorsal attention networks, overlapping the right inferior frontal cortex. The ventral attention network responds with bottom-up inhibition of ongoing action (Aron, Robbins, & Poldrack, 2004, 2014), such as halting manual or speech movement (Aron & Poldrack, 2006; Aron, 2007; Chevrier et al., 2007; Xue et al., 2008). Correspondingly, damage to prefrontal regions affects the ability one has in halting action in response to a stop signal (Aron et al., 2003), and is similarly diminished when the pars opercularis is deactivated with TMS (Chambers et al., 2006). The salience response may also engage the dorsal attention network to facilitate a top-down response (Dosenbach et al., 2007; Eckert et al., 2009; Corbetta & Shulman, 2002; Fox et al., 2006), for example, in goal-directed vocal compensation to pitch-shift (Riecker et al., 2000; Zarate and Zatorre, 2005; Toyomura et al., 2007) or somatosensory perturbation (Golfinopoulos et al. 2011). The localization of the right IFG ROI in the current study was determined by an ALE meta-analysis on neuroimaging studies that experimentally manipulated auditory feedback from both vocal and manual production (Johnson et al., 2019). As the current experimental design required no explicit response to a change in stimulus quality, we hypothesized that increased activity in the right IFG pars opercularis may represent the initial salience response to unexpected voice quality. However, the effect of voice identity in the right IFG did not reach significance, and there was no significant interaction between stimulus source and voice identity in this region. We note that the main effect of source appears most strongly driven by unfamiliar or ambiguous voices, with an intermediate level increase in the uncertain condition (see Figure 4B). It is possible that substantial variability in the data limiting these results was due to the passive nature of the task with no overt attention to the stimulus quality. As activity in this region is associated with attention and subsequent inhibition/adaptation responses, the degree to which each participant attended to the change in stimulus quality is unclear. Furthemore, although psychometric testing confirmed the subjective ability of participants to correctly recognize voice as their own or another speaker’s at a behaviour level, it is possible that the brief vowel stimuli did not provide sufficient information to signal a strong response to unexpected changes in self-voice. Further research is therefore needed to clarify whether the right IFG is responsive to voice identity, and to which extent this may be driven by the degree of salience elicited in divergence from expected qualities of self-voice.

### VARIABILITY IN SELF-MONITORING THRESHOLDS

Although recordings of self-voice can produce a feeling of eeriness for listeners as compared to when spoken (Kimura et al., 2018), people nevertheless recognize recorded voice samples as their own (Nakamura et al., 2001; Kaplan et al., 2008; Rosa et al., 2008; Hughes & Nicholson, 2010; Xu et al., 2013; Candini et al., 2014; Pinheiro et al., 2016a, 2016b, 2019). However, in ambiguous conditions (i.e. acoustic distortion), the ability to accurately attribute a voice to oneself becomes diminished (Allen et al., 2004, 2005, 2006, 2007; Fu et al., 2006; Kumari et al., 2010a, 2010b). As ambiguity increases, an attribution threshold is passed, initiating a transition from uncertainty to externalization (Johns et al., 2001, 2003, 2006; Vermissen et al., 2007). This threshold however varies from person to person (Asai and Tanno, 2013). Here, it was therefore necessary to determine the degree of morphing required to elicit uncertainty in the attribution of voice identity via separate 2AFC psychometric analysis for each participant. In doing so, we could confirm that fMRI responses to the PMA condition were specific to the experience of maximum uncertainty, regardless of any variability in the individual thresholds present in our healthy sample. The results confirmed that participants were able to discriminate their self-voice from an unfamiliar voice, with relatively little variation regarding the point of maximum ambiguity.

In contrast, it is known that persons with schizophrenia display a bias to misattribute self-voice to an external source, both when they listen to recordings of their voice (Ilankovic et al., 2011; Kambeitz-Ilankovic et al., 2013) and when they are speaking (Kumari et al., 2008, 2010b; Sapara et al., 2015). This externalization bias is particularly prominent in schizophrenia patients who experience AVH (Johns et al., 2001, 2006; Allen et al., 2004, 2007; Heinks-Maldonado et al., 2007; Costafreda et al., 2008). Moreover, these individuals are highly confident in their misattributions, as they are more likely to perceive a voice in ambiguous conditions as external rather than remaining uncertain (Johns et al., 2001; Allen et al., 2004; Pinheiro et al., 2016a). It was hypothesised that voice misattribution may underlie AVH as self-voice, either spoken aloud or subvocalized, is mistaken for the voice of an external agent (Frith & Done, 1988; Bentall, 1990; Brookwell, Bentall, & Varese, 2013). Correspondingly, as the severity of AVH symptoms increase, accuracy in self-attribution voice diminishes (Allen et al., 2004, 2006; Pinheiro et al., 2016a). Furthermore, the propensity to externalize self-voice has been linked to hypersalient processing of auditory signals seen in persons with schizophrenia and other populations experiencing AVH (Waters et al., 2012). Notably, this symptomology does not only exist within patient groups. Individuals who are sub-clinical but at a high risk to develop psychosis, display levels of self-monitoring performance similar to patients who meet a clinical diagnosis of schizophrenia (Vermissen et al., 2007; Johns et al., 2010). Indeed, proneness to hallucinate is a continuum and AVH are experienced in the general populations as well, although at lower rates (Baumeister et al., 2017). Even in non-clinical populations, AVH are associated with a bias towards external voice attributions (Asai & Tanno, 2013; Pinheiro et al., 2019). The current findings may be of value in the understanding of the neural substrates underlying dysfunctional self-other voice attribution. In light of our observation that the aSTG displays a qualitatively different activation tendencies for self-voice relative to an unfamiliar voice and the hypothesized influence of right IFG overactivity in salience detection in AVH, we suggest future research in high risk groups to assess a possible abnormal interaction between these two regions. Structural and functional connectivity MRI analysis may help explain if it is abnormalities in the communication between these two regions, or individual disturbances in either or both regions that leads to this symptomatology.

## 5. CONCLUSION

The goal of this experiment was to investigate how levels of self-voice certainty alter brain activity in voice identity and feedback quality monitoring regions of the brain. By replicating earlier findings using a voice area localizer task, we isolated a putative voice identity processing region in the right aSTG. Our results indicate activity in this TVA is suppressed only when self-generating a voice that is definitively attributed to oneself. Furthermore, in the right IFG pars opercularis region responsive to unexpected feedback quality, we demonstrate increased activity while monitoring voice during action relative to when passively heard. It is possible that this activity is driven by salience responses to self-produced stimuli that do not match the expected quality of self-voice. Using a novel self-monitoring paradigm, we provide the first fMRI evidence for the effectiveness of button-press voice-elicitation in modulating an identity-related MIS in the auditory cortex. Furthermore, we present novel findings on the effectiveness of brief vowel excerpts to provide sufficient paralinguistic information to explicitly identify one’s own voice identity. Finally, we suggest a dynamic interaction between the roles of the right aSTG and IFG in the voice self-monitoring network. One may speculate that the feedback monitoring frontal region informs the temporal identity region whenever a salience threshold has been passed and voice feedback is influenced by or under control of an external actor. The implications of variability in the function of these mechanisms are particularly relevant to AVH and may provide specific substrates for the symptomatology seen across the population, independent from broader neural dysfunction associated with clinical pathology.

## ACKNOWLEDGMENTS

This work has been supported by the Fundação Bial, Grant/Award Number: BIAL 238/16; Fundação para a Ciência e a Tecnologia, Grant/Award Number: PTDC/MHC-PCN/0101/2014. Further funding was provided by the Maastricht Brain Imaging Center, MBIC Funding Number: F8000E14, F8000F14, F8042, F8051. We thank Lisa Goller for support in coordination and data collection.

## COMPETING INTERESTS

All authors disclose no potential sources of conflict of interest.

## AUTHOR CONTRIBUTIONS

JFJ, MB, MS, APP, & SAK designed the experiment. JFJ collected the data. JFJ analysed the data with methodological feedback from MB, MS, and SAK. JFJ wrote the manuscript and MB, MS, APP and SAK provided feedback and edits. APP, MS, SAK secured funding.

## Notes

### Competing Interest Statement

The authors have declared no competing interest.

